# Autoregulation dependent and independent mechanisms are responsible for the systemic control of nodule formation by the plant N demand

**DOI:** 10.1101/2021.02.03.429583

**Authors:** Marjorie Pervent, Ilana Lambert, Marc Tauzin, Alicia Karouani, Martha Nigg, Marie-Françoise Jardinaud, Dany Severac, Stefano Colella, Marie-Laure Martin-Magniette, Marc Lepetit

## Abstract

In legumes interacting with rhizobia the formation of symbiotic organs responsible for the acquisition of atmospheric nitrogen is depending of the plant nitrogen (N) demand. We discriminated between local and systemic impact of nitrogen on nodule formation using *Medicago truncatula* plants cultivated in split-root systems. We obtained evidence of the control of nodule formation by whole plant systemic N-satisfaction signaling but obtained little evidence of a local control by mineral nitrogen. We characterized the impact of systemic N signaling on the root transcriptome reprogramming associated to nodule formation. We identified, large genes clusters displaying common expression profiles in response to systemic N signaling enriched in particular fonctions required during these biological processes. We found evidence of a strong effect of SUNN in the control by systemic N signaling of many genes involved in the early interaction with rhizobium as well as organogenesis supporting a role of autoregulation pathway in systemic N signaling. However, we also found evidence that major SUNN independent systemic N signaling controls were maintained in the mutant. This study shed light on the unexpected high complexity of the control of nodule formation by systemic N signaling, that probably involves multiple pathways.

## Introduction

Soil mineral nitrogen (N) availability is a major limiting factor of plant growth and crop productivity. Legume holobionts associated with rhizobia may escape from mineral N limitation because of their unique capacity to acquire the unlimited N source of atmospheric N_2_. The symbiotic interaction of N-limited legume roots with compatible rhizobia results in the formation of organs called nodules generally formed in roots. In the symbiotic organs, the nitrogenase, expressed in differentiated bacteroids, is responsible for the reduction of N_2_ to NH_4_^+^, subsequently exported into the cytosol of the infected plant cells. However, symbiotic N fixation (SNF) is functionally highly dependent on the plant (Oldroyd *et al*., 2011). The allocation of sucrose from the shoot to the symbiotic organs is the source of carbon and energy fueling the bacteroids. Ammonium assimilation by bacteroids is repressed, making them dependent on amino acids supplied by the plant (Prell *et al*., 2009). Leghemoglobins expression by the plant allows the low-oxygen environment required for the nitrogenase activity in bacteroids (Ott *et al*., 2005; Larrainzar *et al*., 2020). In the last decades, mechanisms behind infection of roots by rhizobia and nodule formation began to be unraveled, notably in model legumes *Medicago truncatula* and *Lotus Japonicus* (Oldroyd *et al*., 2011; Mergaert *et al*., 2020). Secretion by the bacteria of lipo-oligosaccharide nod factors and recognition by the plant of compatible bacteria allow the infection and the activation of a complex pathway that results finally in the development of the symbiotic organs. An extensive transcriptome reprogramming is associated with this process (review by Mergaert *et al*., 2020). It involves early and late up-regulation of hundreds of genes involved in early signaling responses, bacterial infection, organogenesis, rhizobium colonization and differentiation, and SNF activation in the roots. A network of plant hormones also contributes to the symbiotic developmental program (Ferguson and Mathesius, 2014; Buhian and Bensmihen, 2018). Ethylene, cytokinins and auxins have been implicated at different stages of infection and nodule formation (Reid *et al*., 2011*b*; Guinel, 2015; Gamas *et al*., 2017).

Studies in many plant species (including legumes not associated with their symbionts) showed that NO_3_^-^ acts locally as a signal stimulating root development and NO_3_^-^ acquisition and assimilation. These regulations contribute to root NO_3_^-^ foraging enabling the sessile plant to preferentially explore NO_3_^-^ rich soil patches (Drew, 1975; Forde and Lorenzo, 2001; Li *et al*. 2014; Gent and Forde, 2017). Genes involved in NO_3_ ^-^ sensing and the large transcriptome and hormonal reprogramming associated with the response to NO_3_^-^ were discovered (Krouk *et al*., 2010; Vidal *et al*., 2020). However, mineral N also assimilates into downstream N metabolites required for whole-plant growth. The integration of N nutrition at the whole plant level leads to adjusting the NO_3_^-^ acquisition capacities to the whole plant N demand. The use of split-root systems experimentally revealed inter-organ systemic signaling controlling root N acquisition (Gansel *et al*., 2001). Molecular mechanisms behind these systemic regulations begin to be unraveled (Okamoto *et al*., 2016; Ohkubo *et al*., 2017; Ota *et al*., 2020). Another whole plant control concerns the regulation by photosynthetates of transporter genes expressed in the roots allowing the root N acquisition to match the shoot photosynthetic capacity (Lejay *et al*., 2003, 2008; Chaput *et al*., 2020). Although these local and systemic signaling mechanisms can be discriminated conceptually and experimentally, it is a major challenge to decipher their interactions and crosstalks because they act synergistically, they share many targets and they contribute to pleiotropic aspects of plant physiology. For example, the three signaling pathways regulate the major root high-affinity NO_3_^-^ transporter AtNRT2.1 involved in NO_3_^-^ uptake in *Arabidopsis* and are interacting with many aspects of plant physiology (Chaput *et al*., 2020). The N status of the plant strongly determines also symbiotic organ development and functioning. Successful nodule formation depends on the whole plant N limitation (Jeudy et al., 2010), and nodule senescence is activated when the holobiont has a sufficient mineral N supply (Pérez Guerra *et al*., 2010). The N signaling mechanisms controlling symbiosis are not as well characterized as those involved in the control of mineral N acquisition. Both high level of NO_3_^-^ supply and accumulation of downstream N metabolites repress symbiosis activity (Silsbury *et al*., 1986; Bacanamwo and Harper, 1997; Ruffel *et al*., 2008). Nevertheless, in *Lotus*, low levels of nodule NO_3_^-^ intake stimulates nodule functioning (Valkov *et al*., 2017). There is also evidence of local suppression by the plant of symbiosis in nodules that do not fix N_2_ that was qualified as a “sanction” mechanism against ineffective symbiotic partners (Kiers *et al*., 2003). Major impact on symbiosis of systemic N signaling was reported, particularly in *Medicago truncatula* (Jeudy *et al*., 2010; Laguerre *et al*., 2012; Lambert *et al*., 2020*b*). The provision of ample levels of mineral N to symbiotic plants induces a systemic signal that activates rapidly nodule senescence (Lambert *et al*., 2020*b*). The partial suppression of SNF by Ar/O_2_ localized treatment in split-root systems systemically stimulates pre-existing nodule expansion and nodule formation to compensate for the N-deficit of the plant (Jeudy *et al*., 2010; Laguerre *et al*., 2012; Lambert *et al*., 2020*b*). Systemic responses to variations of the symbiotic plant N demand correlates to variations of shoot-root sucrose translocation and/or hormonal pools (Lambert *et al*., 2020*b*). There are contrasting reports about the balance between local and systemic regulations of nodule formation by mineral N. Localized repression of nodule formation by NO_3_^-^ was reported in soybean (Hinson, 1975; Tanaka *et al*., 1985; Cho and Harper, 1991; Xia *et al*., 2017), but the systemic repression of nodulation by mineral nitrogen supply seems to be proeminent in *Medicago truncatula* (Jeudy *et al*., 2010; Kassaw *et al*., 2015).

In legume holobionts the autoregulation mechanism (AON) is involved in the systemic control of nodulation. It enable the earliest formed nodules to suppress further nodulation (Kosslak and Bohlool, 1984). Because AON activates at the early stages of the interaction during nodule development, far before SNF become active, AON cannot be considered as a feedback mechanism related to the satisfaction of the N demand and/or the accumulation of downstream N metabolites (Kosslak and Bohlool, 1984; van Brussel *et al*., 2002; Li *et al*., 2009). However, several lines of evidence argue for a role of AON in N signaling. AON mutants form high number of nodules (“hypernodulating” mutants) under a high mineral N supply that is a normally suppressive condition for nodulation in wild types (review by Reid *et al*., 2011*b*). The suppression of SNF by Ar/O_2_ treatments on split-root plants revealed that whole-plant N-limitation releases the systemic inhibition of nodule formation by autoregulation (Jeudy *et al*., 2010; Laguerre *et al*., 2012). This mechanism involves a leucine-rich repeat receptor-like kinase (LRR-RLK, reviewed by Mortier *et al*., 2012*b*) identified in several legume species. In *Medicago truncatula* it is encoded by *SUNN* (Schnabel *et al*., 2005). CLE peptides produced in the roots in response to the interaction with *rhizobium* are translocated from the root to the shoot, where they associate with this receptor resulting to a systemic inhibition of nodule formation (Mortier *et al*., 2012*a*; Okamoto *et al*., 2013). This inhibition is associated, both in *Lotus* and *Medicago*, with a lower translocation from the shoot to the root of the miRNA miR2111 (Tsikou *et al*., 2018; Nishida and Suzaki, 2018; Gautrat *et al*., 2020). Shoot cytokinin (CK) and methyl jasmonate accumulation might also have a role in the systemic control of nodulation in *Lotus japonicus* (Nakagawa and Kawaguchi, 2006; Kinkema and Gresshoff, 2008; Sasaki *et al*., 2014; Azarakhsh *et al*., 2018). In soybean, *Lotus*, and *Medicago*, some CLE peptides, able to activate AON, accumulate in roots in response to NO_3_^-^ through a pathway that may implicate NIN and other NLP transcription factors (Reid *et al*., 2011*a*; Nishida *et al*., 2018; Mens *et al*., 2020). Unexpectedly, these observations suggest an activation of systemic repression of nodulation resulting from local sensing of NO_3_^-^ rather than downstream N metabolites accumulation. Increasing evidence suggest that both AON dependent and independent systemic signaling mechanisms control nodule development (Jeudy *et al*., 2010; Kassaw *et al*., 2015). The nodule expansion response to systemic signaling of plant N deficit remains active in the sunn mutant (Jeudy *et al*., 2010). A systemic mechanism controlling nodule formation implicating MtCRA2, another LRR-RLK acting in shoots in *Medicago truncatula*, positively regulates nodule formation in parallel with the classical SUNN/AON pathway (Huault *et al*., 2014; Laffont *et al*., 2019, 2020). MtCRA2 is activated by small peptides of the CEP family produced in roots exposed to mineral N deprivation or rhizobium (Mohd-Radzman *et al*., 2016). In *Arabidopsis*, CEPR1, the homolog of CRA2 was found to also interact with CEP peptides and mediate a systemic signaling pathway responsible for adjusting NO_3_^-^ uptake capacity to the plant N demand (Tabata *et al*., 2014; Ota *et al*., 2020). However, up to date, the CRA2/CEP pathway’s contribution to the regulation of symbiosis by N signaling remain poorly understood.

The molecular targets of systemic N signaling controlling nodule formation remains poorly characterized. A few studies have shown that the addition of mineral N to the roots of legume-rhizobium holobionts is associated with nodule transcriptome reprogramming (Omrane *et al*., 2009; Moreau *et al*., 2011; Seabra *et al*., 2012; Cabeza *et al*., 2014). However, these studies did not discriminate between the local effects of mineral N (i.e., at the site of application) and the systemic effects (i.e., related to the satisfaction of the whole-plant N demand). A previous report based on split-root systems analysis has shown that whole-plant systemic N signaling has a substantial impact on the transcriptome of nodulated roots, but the effects on nodule formation and/or mature nodule were not separated (Ruffel *et al*., 2008). In a recent study, we characterized the impact of systemic N signaling on the mature nodules in the *Medicago truncatula*/*Sinorhizobium medicae* holobiont (Lambert *et al*., 2020*b*). In the present study, we characterized the control of nodule formation by systemic N signaling and identified the root transcriptome response to systemic N signaling using RNAseq in the same biological model. The *Medicago truncatula sunn* mutant was compared to the wild type to investigate the contribution of AON to this control.

## Results

### Whole plant N signaling controls nodule formation

We characterized the effect of plant’s N status on nodule formation using split root systems on *Medicago truncatula A17* inoculated with *Sinorhizobium medicae md4* (Fig.1A). Plants were cultivated hydroponically. We separated roots of individual plants into two compartments and we applied contrasted nutrient regimes. We supplied one half of the plant’s root system with high level (10 mM NH_4_NO_3_; SNO) or low level (0,5 mM KNO_3_; LNO) of mineral N resulting respectively in N-satisfied or N-limited plants. We provided an aerated nutrient solution without mineral N to the second half root systems of these plants (SN and LN, respectively). We inoculated the roots of all the split-root compartments with *Sm* md4 two days after establishing the N-treatments. As expected, these nutrient regimes resulted in higher shoot dry weight in N-satisfied plants than in N-limited plants (Supplementary Table S1). We compared half root systems of the same plant to estimate the local effect of the N treatments. Adding NO_3_^-^ (high or low concentrations) stimulated root proliferation at the application site doubling the root length normalized per the shoot biomass (NRL; Fig. 1B, Supplementary Table S1, comparison SNO vs SN and LNO vs LN). Comparing roots exposed to the same local environment but belonging to N-satisfied or N-limited plants allowed investigating the whole plant systemic N signaling. The repression of root development by systemic N-satiety signaling resulted in a NRL reduction in N-satisfied plants compared to N-limited plants (Fig. 1B, Supplementary Table S1, comparison SN vs LN). Nodulation was repressed in N-satisfied plants as compared to N-limited plants (Fig. 1C, Supplementary Table S1). The repression of nodule formation in SN roots (indirectly exposed to the treatment) resulted from a systemic N satisfaction signaling (Fig. 1C, Supplementary Table S1, comparison SN vs LN). Local effects of mineral N availability on root nodule density were also evidenced (Fig. 1C, Supplementary Table S1, comparison SNO vs SN). However, because the relative proportion of nodules in the treated and untreated half root systems of N-satisfied or N-limited plants were equivalent (Fig. 1D, Supplementary Table S1), these nodule density differences were more likely explained by the stimulation of root growth rather than by a direct effect on nodulation. Interestingly, although root and nodule developments were controlled, both by local and systemic signaling, their responses to N treatments were rather different. As expected, root development (root length) was mainly resulting of local signaling of mineral N presence, whereas nodule formation was mainly resulting of whole plant N systemic signaling. We further confirmed the morphological analysis by comparing LN and SN responses to inoculation in stable transgenic plants expressing *pENOD11:: GUS* reporter gene cultivated in split root systems of Fig1A. Both SN and LN roots displayed early infection responses at 1 and 2 days post-inoculation (dpi) that were more attenuated in SN compared to LN at 4 dpi (Supplementary Fig. S1).

**Fig. 1.**
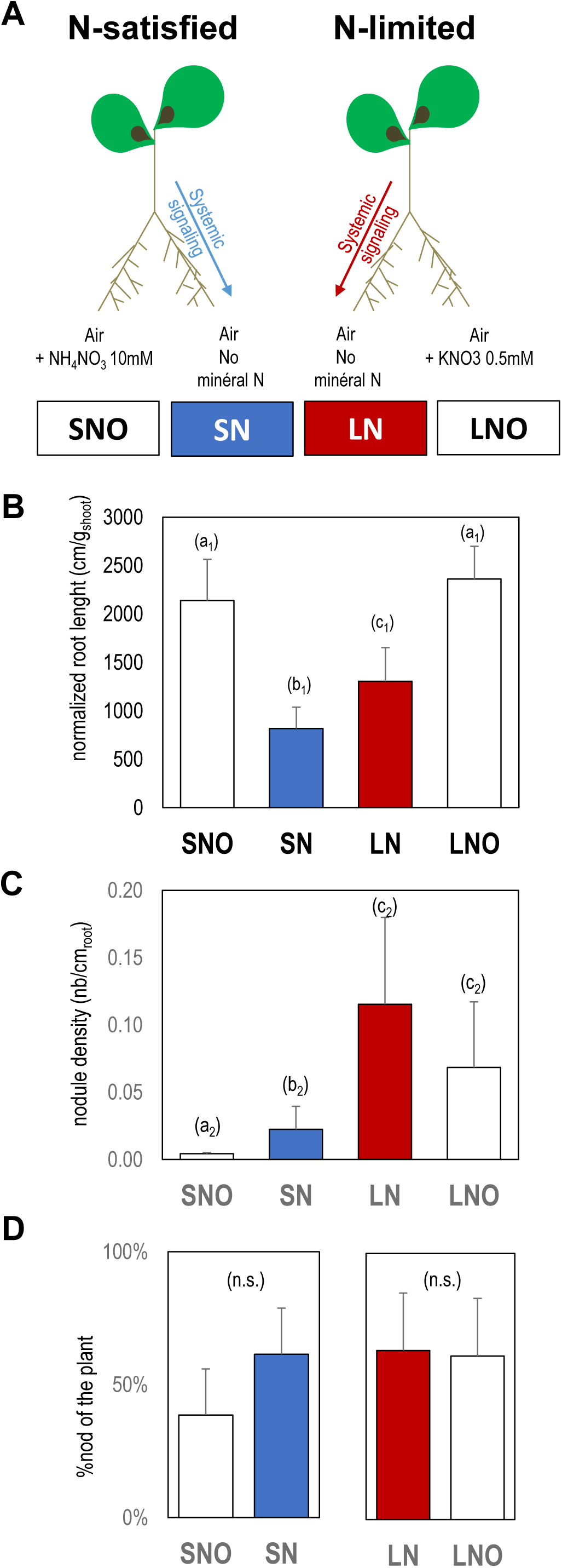
Split-root systems used to study the response of nodule formation to N-satisfaction and N-limitation signaling. A. Plants were cultivated hydroponically for 6 weeks. N-treatments were applied on one half of the root systems (SNO, LNO) 12 days before harvest. Effects of the treatments were studied on the other sides of the root systems supplied with nutrient solution without mineral nitrogen (SN in blue, LN in red). N satisfaction or N limitation were achieved by providing 10 mM NH_4_NO_3_ or 0,5 mM KNO_3_ respectively. All roots were inoculated with *Sinorhizobium medicae* md4 48h after initiating the N treatments. **B**. Normalized root length per shoot dry mass (NRL). **C**. Nodule density (nodule number per root length). **D**. Proportion of nodules (%) present in both compartments of the split-root systems. Detailed data are provided in <u>Supplemental Table S1</u>. Letters indicate distinct groups of values deduced from ANOVA and pairwise t-test using a p-value threshold of 0.05. n.s. indicates a non significant difference. Values are means±SD (n=5).

### Impact of N signaling on the transcript reprogramming associated with plant-*rhizobium* interaction

Firstly, the transcriptome reprogramming associated the plant-rhizobium interaction already described by previous studies was confirmed in our split-root system. We investigated the effect of the Sinorhizobium inoculation (24h, 48h) on the LN roots of N-limited plants (Fig. 1A). A total of 11464 transcript responsive to the inoculation (RRTs) were identified (Supplementary Table S2). As expected, they included typical transcripts encoding early nodulins, transcriptions factors, structural or regulatory proteins already characterized as markers of the induction of the rhizobium infection and nodule organogenesis programs (Supplementary Table S3). The activation of these genes correlates to the downregulation of transcripts encoding transporters and enzymes involved in the NO_3_^-^ utilization (Supplementary Table S3).

Using a statistical modeling of the RNAseq data, we investigated the effect of the systemic signaling of the plant N status on the transcriptome by comparing the LN roots of N-limited plants to the SN roots of N-satisfied plants at 2, 4, and 7 dpi. Differential analyses at the different inoculation times (Fig.2) indicated that the number of transcripts differentially accumulated in LN vs SN roots increased strongly on the 4-7 days period as compared to the 2-4 days period. This observation suggested that the responses to systemic N-signaling of roots inoculated by rhizobium varied during the interaction. A total of 8133 N-responsive differentially accumulated transcripts (N-resp DATs) in at least one of these pairwise comparisons (2,4 or 7 dpi) were identified (Supplementary Table S4). Only 25% of the RRTs are N-resp DATs, which was a highly significant but a marginal fraction. However, RRTs were particularly abundant in the N-resp DATs (57%) indicating that most of the roots transcripts regulated by systemic N signaling responded also to rhizobium (Fig.3). Globally, the N-resp DATs were significantly enriched in transcripts of genes belonging to symbiosis related islands (SRI) of the medicago genome (36% of N-resp DATs) described by Pecrix *et al*. (2018) as well as in transcripts specifically accumulated in the nodule (35% of the N-resp DATs) described by Roux *et al*. (2014). Many transcripts associated with the late phases of nodule organogenesis and SNF maturation belonged to N-resp DATs (Fig.3). A strong impact of systemic signaling was particularly noticed on transcripts associated with bacteroid differentiation. The accumulation of numerous transcripts encoding NCR and GRP peptides families (Fig. S2) as well as MtDNF1, MtDNF2, and MtCCS52a (Fig.3) strongly depended on systemic N signaling (i.e. up-regulated in N-limited plants as compared to N-satisfied plants), as well as transcripts involved in nodule functioning such as leghemoglobin and the sugar efflux transporter MtSWEET11 potentially involved in nodule sugar allocation.

**Fig. 2.**
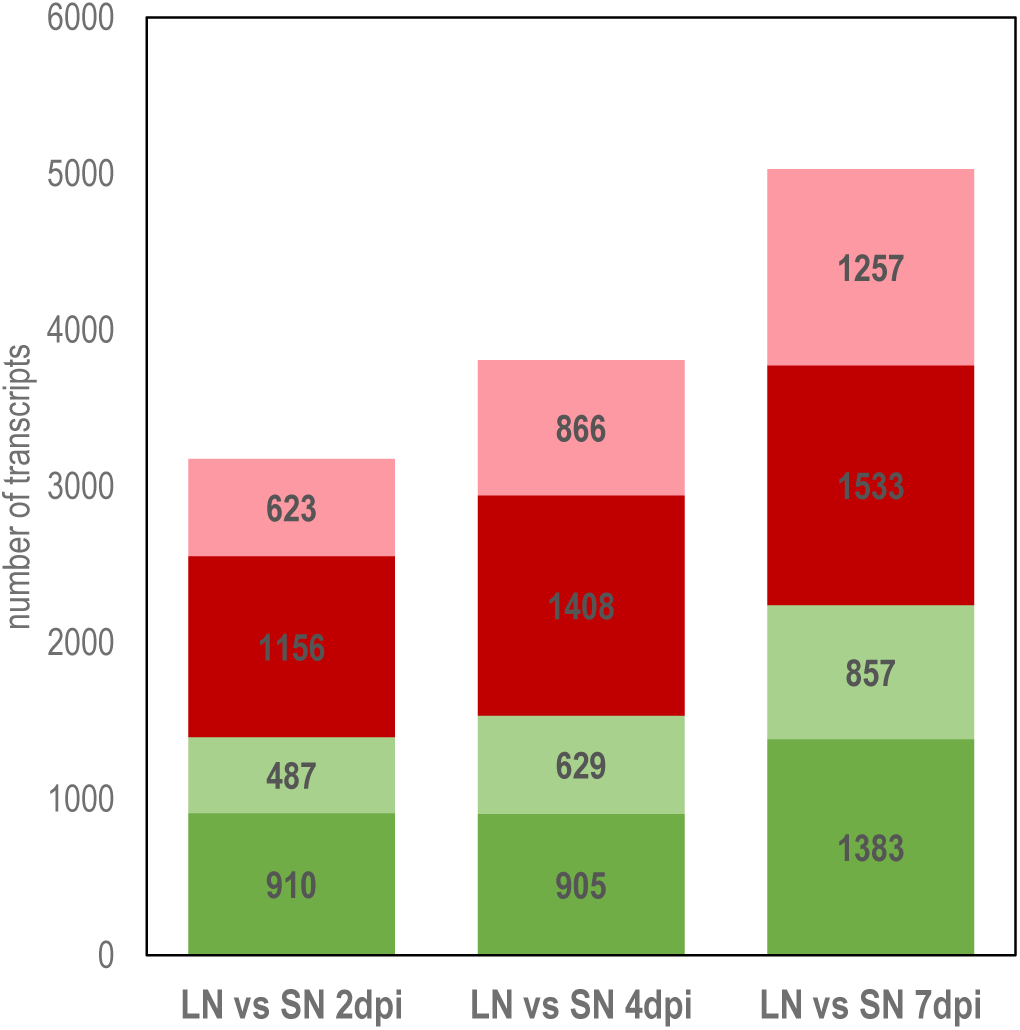
Effect of systemic N signaling on the transcriptome of inoculated root. LN and SN roots were compared at 2, 4 and 7 dpi. Red and green area represent respectively transcripts over-accumulated and under-accumulated in DN roots as compared to SN roots. Dark colors represent RITs.

**Fig. 3.**
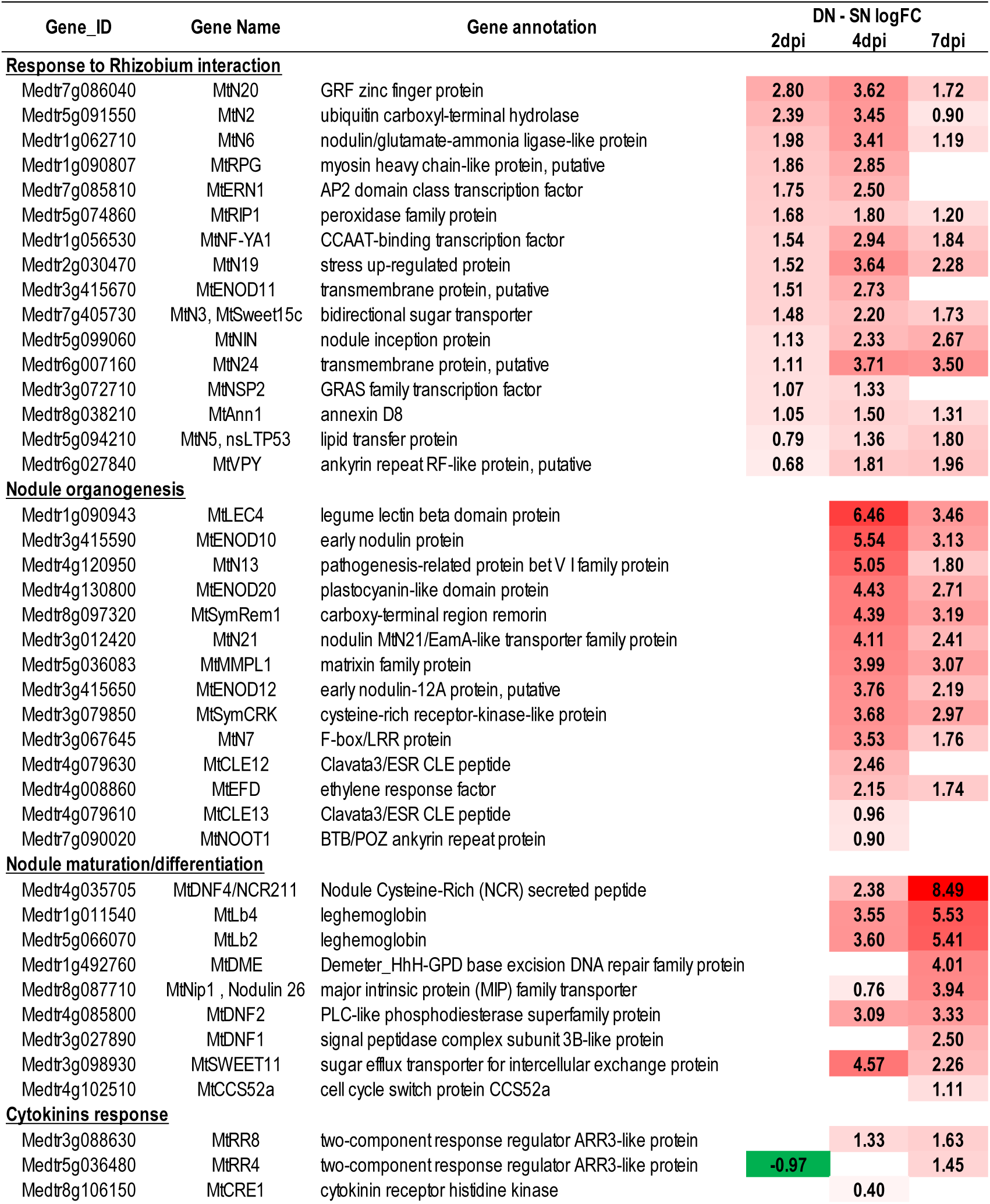
Heat map of the effects of systemic N signaling on the expression of marker transcripts associated to nodule formation by previous studies. LN and SN roots were compared at 2, 4 and 7 dpi (supplementary table 3). Only significant FC difference (p-value and FDR <0.05) are indicated. Red and green colors represent transcripts respectively up-regulated and down regulated in LN droots as compared to SN roots.

The co-expression analysis based on mixture models organized the N-resp DATs in 10 clusters according to their accumulation kinetics in LN and SN inoculated roots (Fig.4A; Supplementary Table S5). The model fitted well the data as only 10% of transcripts were not classified. The 10 clusters can be grouped according to the transcriptional profile in response to N-limitation signaling: up-regulation in clusters 1, 2, 4, 5, 8, 10, and down-regulation in clusters 3, 6, 7, 9 (Fig. 4A). As expected, RRTs identified in our initial response to inoculation analysis (1 and 2 dpi) were highly represented in all these clusters except for cluster 4 that gathered only transcripts activated in the late phases of nodule formation: bacteroid differentiation (NCRs, GRPs), activation of SNF (Leghemoglobins). Compared to the the whole annotated genome, each cluster is associated with specific functions, as shown in Fig.4B (p<0,05 hypergeometric test ; Supplementary Table S5)). Cluster 3 was particularly enriched in transcripts related to NO _3_^-^ utilization, while Clusters 5 and 4 gathered many “nodulins” transcripts. Clusters 2 and 4 contained most of the NCRs, GRPs peptides, and leghemoglobin transcripts, as well as the sugar transporter MtSWEET11 transcript

**Fig. 4.**
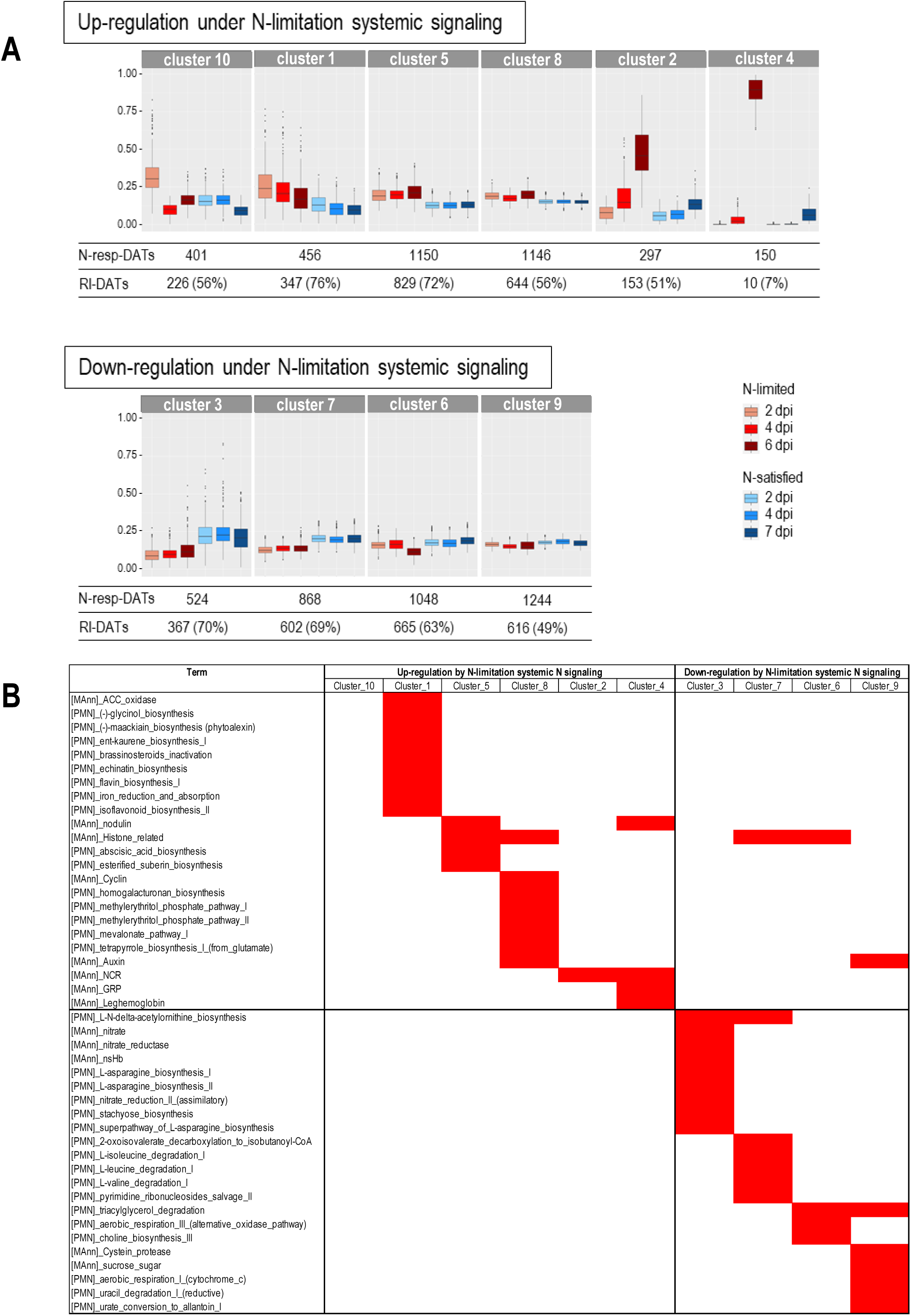
Co-expression analysis of the transcripts differentially accumulated in response to systemic N signaling in rhizobium inoculated roots. LN and SN roots were compared at 2, 4 and 7 dpi. N-resp DATs are listed in Supplementary Table S3. A. Co expression clusters and their normalized accumulation kinetics (Coseq package). B. Specific annotation enrichment of the cluster in specific functions as compared to entire annotated genome (estimated by hypergeometric test).

### Role of AON in the repression of nodule formation by systemic N satiety signaling

We compared the *sunn* mutant impaired in the LRR-RLK required for AON signaling in *Medicago truncatula* and WT A17 plants in split-root systems (Fig. 1). We monitored the expression of 6 marker transcripts known to be up-regulated at various stages of the nodulation process (Fig. 5). Transcripts encoding the early nodulin MtENOD11 (Journet *et al*., 2001) or transcription factors NIN (Marsh *et al*., 2007), NFY-A1 (Combier *et al*., 2007), NSP2 (Kaló *et al*., 2005) and ERN1 (Middleton *et al*., 2007) orchestrating transcriptome reprogramming associated to rhizobium-legume interaction mark the early infection. MtRR4 (Gonzalez-Rizzo *et al*., 2006), involved in the cytokinin response, or MtMMPL1 (Combier *et al*., 2007), involved in the progression of the infection, associate with nodule organogenesis. Late stages of nodule formation are marked by the up-regulation of large numbers of transcripts encoding nodule-specific cysteine-rich (NCR) associated with bacteroid differentiation (Kereszt et al., 2018) as well as the genes encoding leghemoglobins allowing the bacterial nitrogenase to be active in a microoxic environment (Ott *et al*., 2005).Transcripts levels were quantified by RT-qPCR before inoculation and 1, 2, 3, and 7 dpi in LN and SN root systems. In parallel, we monitored the activity of *pENOD11::GUS* reporter gene in roots of stable transgenic plants of the same genotypes (Supplementary Fig. S1). As expected the 6 transcripts were all up-regulated in response to *Sinorhizobium* in LN roots. They were all regulated by systemic N signaling in A17. *MtENOD11, MtNIN*, and *MtNFYA1* were up-regulated rapidly at the early stages of the interaction in both SN and LN roots. Their responses to *Sinorhizobium* at 3-7 dpi were attenuated in SN roots compared to LN roots. *MtRR4, MtMMPL1, MtNCR084*, and *Leghemoglobin* were activated in LN roots only after 2 dpi (Fig.5), and their activations were reduced in SN roots as compared to LN roots. Both RT-qPCR and *pENOD11::GUS* reporter gene analysis support that *sunn* mutation prevented the repression by N satiety of MtNIN, MtNFYA1, MtENOD11, MtRR4, MtNCR084 transcripts (Fig. 5 and Supplementary Fig.S1). However, this was not true for the leghemoglobin that remained repressed by N-satisfaction signaling in both *sunn* and A17. Although the *sunn* mutation partially released the repression of nodulation by N satisfaction systemic signaling, a *sunn*-independent systemic repression by N satisfaction remained active on the leghemoglobin transcript.

**Fig. 5.**
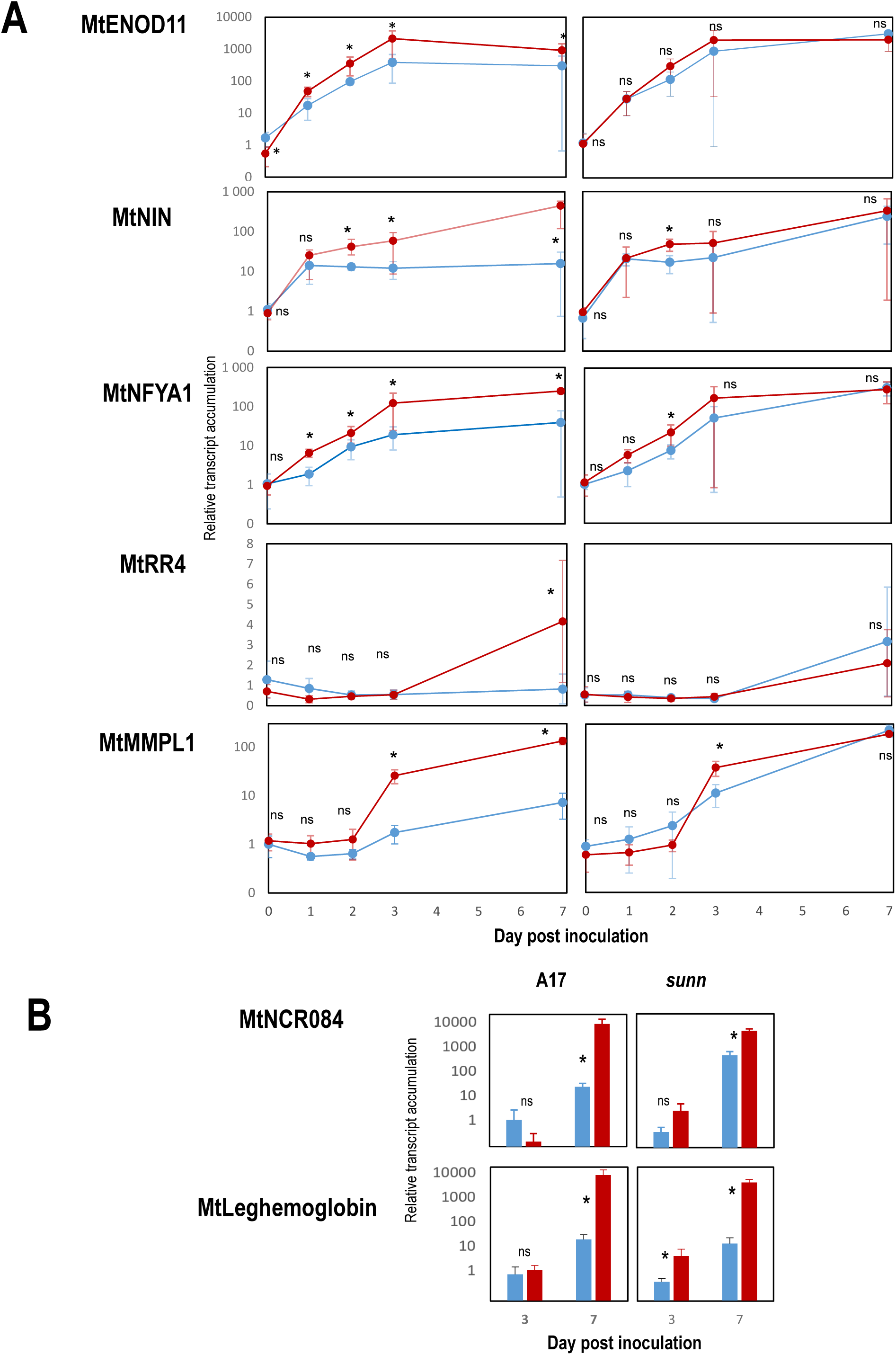
Accumulation of selected transcripts in rhizobium-inoculated roots under contrasted systemic N-signaling. Plant were cultivated in the split root systems described in Fig1.A. Inoculated LN (in red) and SN (in blue) roots belonging to respectively to N-limited and N-satisfied plants were compared at 0, 1, 2, 3 and 7 dpi. Total RNA was extracted from the roots and transcript accumulation was quantified by RT-Q-PCR. A. Kinetic of of MtENOD11, NFYA1, NIN, MtRR4, MtMMPL1 transcripts. B. Transcript accumulation of MtNCR084 and MtLeghemoglobin at 3 and 7 dpi (transcripts not detected at 0, 1 and 2 dpi). Values are means +/-SD, n=6. At each time point LN and SN values were compared according t-test (0.05). * significant difference. ns. non significant difference.

We extended these observations to the whole genome by comparing the A17 and *sunn* responses of the rhizobium inoculated root transcriptome to systemic N signaling (SN vs LN) at 2 and 7 dpi. Transcripts displaying either differential or equivalent responses to systemic N signaling in both genotypes were discriminated by statistical modeling and likelihood ratio tests (Supplementary table S6). The proportion of transcripts displaying a differential response to systemic signaling comparing *sunn* and A17 were higher at 7dpi (43%, 3666/8442) than at 2dpi (11%, 388/3458). The *sunn* mutation impact on the systemic N signaling was stronger during organogenesis and late phases of nodule formation than during the early response to rhizobium. N-resp-DATs were particularly abundant within these transcripts: 60% of transcripts displaying a differential response to systemic signaling between A17 and sunn at 7dpi belong to N-resp-DATs. The overlap was large with N-resp-DATs up-regulated in response to N-limitation (as compared to N satisfaction), particularly for the co-expression clusters 2 and 4 (>90% of the transcripts differentially regulated in sunn and A17). The regulation by N-systemic signaling of typical transcripts known to be associated with rhizobium interaction and nodule formation was clearly impaired in the mutant consistently with its phenotype that results in the maintenance of nodule formation under N-satisfaction signaling (Supplementary Table S7). Nonetheless, our data confirm that not all the responses of the nodulated root to systemic N-satisfaction signaling were impaired in the *sunn* mutant. Most of the transcripts encoding leghemoglobin present in the transcriptome at 7 dpi (8/11) displayed equivalent response to N signaling in sunn and A17 confirming our preliminary observation on a single transcript (Supplementary table S8). Among the 498 NCR transcripts identified in the roots of A17 at 7 dpi, 313 displayed impaired regulation by systemic N signaling in the sunn mutant when compared to A17, but 185 other transcripts displayed equivalent regulation in the two genotypes (Supplementary table S9). Similarly, transcripts encoding GRP peptides or annotated as “nodulins” may be easily discriminated into two categories according to the impact of the sunn mutation on their regulation by systemic N signaling (Supplementary table S10 & S11). This duality of expression profiles did not concern all families of transcripts present at 7 dpi: for example, the systemic N signaling regulation of the 20 transcripts encoding defensin peptides was always depending on *SUNN* (Supplementary table S12). Despite a clear role of *SUNN* in the control of nodule formation by systemic N signaling, altogether, these data demonstrated that additional *SUNN*-independent mechanisms control the late phase of nodule formation/maturation, and may contribute importantly to the adjustment of the symbiotic capacity to the plant N demand.

## Discussion

### Systemic signaling of the whole plant N demand is a major driver of nodule formation

This study provided new insights about the control of the legume-rhizobium by the plant N demand. The success or the abortion of the developmental process initiated by the plant-bacterial interaction are under the control of systemic signaling of the whole plant N demand. This control has a great biological significance. Because symbiotic organ formation and SNF have elevated carbon and energy costs (fulfilled through the allocation of photosynthates by the plant to the symbiotic organs), these whole plant mechanisms allow adjusting the root nodule capacity to the N demand of the entire holobiont. In non symbiotic plants, numerous studies have characterized the local stimulation of root development by NO_3_ ^-^ resulting in root foraging (Drew, 1975; Forde and Lorenzo, 2001; Li et al. 2014; Gent and Forde, 2017). NO_3_^-^ acts as a signal allowing sessile plants to explore and preferentially deplete NO_3_^-^ rich soil patches. This mechanism interacts with a whole plant control that stimulates or represses mineral N uptake and root development as a function of the plant N demand. In symbiotic plants, both root foraging and symbiotic nodule formation mediate plant adaptation to N-limitation (Jeudy et al., 2012), but underlying N signaling processes associated with these two processes are probably different. This study confirms that NO_3_^-^ stimulates the root proliferation locally in inoculated plants equivalently as in non-symbiotic plants (Figure 1B). However, we failed to obtain a clear argument supporting a specific local signaling effect of mineral N on nodule formation. Although we found that NO_3_^-^ reduces locally nodule density (Figure 1C), this may be explained by the stimulation of root expansion (Figure 1B & 1C) rather than by a direct reduction of the nodule formation. We yielded evidences showing that both root development and nodule formation are under the strong control of systemic signaling of the plant N demand (Figure 1B & 1C). Specificities of the underlying developmental processes (nodule vs root development) do not rule out the hypothesis of a common upstream control by a mechanism responsible for sensing and integrating the whole plant N demand. Indeed, mineral N acquisition and SNF fuel the entire plant’s N metabolism and result in the same downstream products. Nevertheless, although many studies have evidenced whole plant N demand regulations and several molecular components associated to systemic N signaling being identified (Li et al., 2014; Okamoto et al., 2016 ; Bellegarde et al., 2017; Jia and von Wirén, 2020), the global underlying whole-plant N sensing mechanism remains elusive.

### Systemic signaling of the whole plant N demand controls the progression of the root transcriptome reprogramming initiated by rhizobium infection

Inter-organ systemic N signaling related to plant N satisfaction and plant N limitation modulates the progression of the transcriptome reprogramming associated with *Sinorhizobium medicae-Medicago truncatula* interaction. Both the number of transcripts differentially accumulated in response to systemic N signaling (Fig. 2) and the amplitude of their responses increased during the plant-rhizobium interaction progression in our experiments (Fig.3&4). The accumulations of many transcripts rapidly up-regulated by rhizobium responded to systemic N signaling, but the impact of systemic N signaling was generally modest at 2 dpi while more pronounced at 4 and 7 dpi (Fig.3). Many typical transcripts that responded rapidly to nod factor signaling, such as, for example, ENOD11, NFY-A1, or NIN, were up-regulated in response to *Sinorhizobium* whatever the N status of the plant (Fig. 5). They were progressively repressed after several days in roots under N-satisfaction signaling but remained activated under N-limitation signaling. Transcripts strongly activated by *Sinorhizobium* at the late stage of the interaction generally responded poorly to rhizobium in roots under N satisfaction signaling. These transcripts were either associated with nodule organogenesis and rhizobium colonization (e.g., MtRR4, MtMMPL1, MtDME), bacteroid differentiation (e.g., NCRs, GRPs), or associated with the activation of SNF (e.g., MtSWEET11, leghemoglobin). We conclude that N systemic signaling does not likely determine the root’s competency to respond to rhizobium but instead controls the symbiotic process leading to progress or abortion of bacterial infection, nodule organogenesis and SNF activation.

### Control of nodule formation by systemic signaling of the plant N demand requires AON-dependent and AON-independent components

AON is frequently interpreted as a negative feedback mechanism allowing the plant to limit symbiotic development to prevent the useless dissipation of energy and photosynthates (Reid *et al*., 2011). Mutants impaired in the SUNN receptor are forming nodules even when supplied with a high level of mineral N, suggesting an important role of AON in the systemic control of nodulation by the plant’s N status (Kinkema et al., 2006). The *sunn* mutation clearly impaired the repression by N satisfaction of numerous transcripts associated with the response to rhizobium, the infection, and the nodule organogenesis in agreement with this hypothesis. However, among the responses associated with systemic N signaling of the rhizobium-legume interaction, the AON pathway constitutes only a part of the puzzle. Firstly, the *sunn* mutation’s impact on the the root transcriptome systemic N signaling responses at 2dpi is significant but marginal. Secondly, although the mutation has a stronger and larger impacts on the N signaling responses of transcripts at 7 dpi, the N-signaling responses of most N-responsive transcripts remain equivalent in the mutant and in the wild type at these late stages of nodule formation. Notably, many nodule specific regulated by systemic N signaling involved in bacteroid differentiation (NCRs and GRPs peptides families), activation of SNF, and sucrose allocation (leghemoglobin, SWEET11) belong to this last group. Our data provided molecular support to previous physiological studies on the *sunn* mutant (Jeudy *et al*., 2010, Kassaw *et al*., 2015), suggesting that the N signaling response of symbiosis had *sunn*-dependent and *sunn*-independent components. They explain why the hypernodulation phenotype paradoxically does not result to a higher level of SNF per plant, despite of a higher nodule biomass per plant (Jeudy *et al*., 2010). Our study provides support for the hypothesis of a *sunn*-independent systemic N signaling mechanism downstream nodule organogenesis controlling SNF activity per nodule. This mechanism has potentially a major role in the adjustment of symbiotic N acquisition capacity to the whole plant N demand. The targets of such mechanism are likely the transcripts encoding proteins involved in bacteroid differentiation, SNF, and sucrose allocation of nodules from the plant. This hypothetic mechanism has features apparently reminiscent of the systemic N-signaling mechanism operating in the mature nodule (Lambert *et al*., 2020b). Wheter they are distinct mechanisms or the same one observed at different stages of nodule development remains to be clarified. A control by the N demand of the allocation by the plant of the C metabolites to the roots fueling the SNF and providing carbon skeleton to amino acid synthesis is consistent with the strong integration C and N metabolisms at the whole plant level (Stitt et al., 2002; Ruffel et al., 2014; Chaput et al., 2020). In addition, this study revealed also a little impact of the *sunn* mutation on the systemic N signaling response observed at 2 dpi, suggesting that a *sunn*-independent mechanism might be also required to explain the N-signaling responses of the early symbiotic interaction. Interestingly another systemic mechanism controling early stage of nodule formation and operating in parallel to the *sunn*-dependent AON have been discovered (Huault *et al*., 2014; Laffont *et al*., 2019, 2020). The possible role of this mechanisms in the systemic N-signaling of early rhizobium-interaction is not known and deserves to be investigated.

### Symbiotic specific or global systemic N signaling mechanisms

Altogether, these data shed light on the high complexity of controls of symbiosis by the plant’s N status and their high level of integration. Because these controls target specific symbiotic processes, they may be interpretated as the result of symbiosis specific mechanisms. However, whether they form specific pathways or are symbiotic components orchestrated by a global signaling hub common to root development and mineral N acquisition, remain to be clarified. There is increasing evidence indicating that SUNN/CLE and CRA2/CEP pathways, initially identified for their role in the systemic control of nodule formation, also have non-symbiotic functions on root development or NO_3_^-^ uptake (Okamoto *et al*., 2013; Tabata *et al*., 2014; Goh *et al*., 2019; Lagunas *et al*., 2019). Furthermore, closely related pathways controlling root development and NO_3_^-^ uptake have been identified in non-legume plants (Tabata *et al*., 2014; Ota *et al*., 2020). This argues for the hypothesis that they have been recruited by symbiosis for a more or less specific function(s) but suggests also they may be components of a global network of systemic N signaling mechanisms controlling and coordinating underground plant development as a function of plant nutritional demand and capacity.

## Materials and methods

### Split-root plant growth condition

The *Medicago truncatula* genotypes were the wild type Jemalong A17 and the TR122 sunn mutant (*sunn*-2 allele ; Sagan *et al*., 1995; Schnabel *et al*., 2005). The transgenic lines carried a *pENOD11 :: GUS* transgene in the A17 (line L416; Charron *et al*., 2004) or in TR122 backgrounds (obtained by genetic introgression of the L416 transgene into sunn). Experimental planning of the split-root experiments is presented in Supplemental Figure S3. Seeds were scarified, germinated as described in Lambert *et al*., 2020. Individual plantlets were transferred into hydroponic culture tanks containing a vigorously aerated HY basal nutrient solution (Lambert *et al*., 2020*b*) adjusted to 5.8 with KOH and supplemented with 1 mM KNO_3_. We cut the primary root tips of plantlets to promote branching of the root system. The culture chambers conditions were a light intensity of 250 μmol s−1 m−2 photosynthetically active radiation, a relative humidity of 70%, a light/dark cycle of 16h/8h and an ambient temperature of 22°C/20°C. We separated the root systems of 4-week-old plants in two parts. Initialy all the split-root compartments contained HY nutrient solution supplemented with 0.5mM KNO_3_ as a low mineral N source and (the pH was adjusted to 7). We initiated the N treatments by providing a HY nutrient solution supplemented respectively with high mineral N (10mM NH_4_NO_3_) and low mineral N sources (0,5mM KNO_3_) to the SNO and LNO compartments whereas the SN and LN compartments contained HY nutrient solution without mineral N (Fig.1A). Roots of all compartments were inoculated 48h after the N treatment initiation with *Sinorhizobium medicae md4* (10^7^ exponentially growing bacteria per ml). We renewed the nutrient solutions every 4 days. We initiated N treatments at different times before harvest in order to compare plants of the same age (6 week old plants) differing by the duration post-inoculation (Supplemental Figure S3). Scanned Image (600 dpi) of the root systems were analyzed by using the Optimas image analysis software (MediaCybernetics) to characterize root growth parameters. GUS histochemical staining procedure was already described (Lagarde *et al*., 1996).

### RNA analysis

The first experiment (exp1) compared the LN A17 roots at 0, 1, 2 day post-inoculation (dpi; 0 is the non-inoculated control). The second experiment (exp2) compared the LN and SN roots at 2, 4 and 7 dpi. The third experiment (exp3) compared the LN and SN roots of A17 and sunn at 2 and 7 dpi. We collected all roots samples simultaneously in LN or SN compartments of the split-root systems. Each biological replicate is a pool of the half root systems of two plants. Total RNA was extracted using QIAzol Lysis Reagent, purified using miRNeasy®, and digested by DNAse I to eliminate DNA contamination according to the supplier’s recommendations (Qiagen). RNAseq analysis included 3 (exp1 and 3) or 4 (exp2) biological replicates per condition. Polyadenylated plant mRNA libraries were generated and sequenced in mode single-read 50 nt on illumina HiSeq 2500 as described previously (Lambert *et al*., 2020*b*). The sequencing reads (ArrayExpress database accession numbers E-MTAB-9932, E-MTAB-9941, E-MTAB-9942) were mapped on the M. truncatula v4.2 using the glint software (T. Faraut and E. Courcelle; http://lipm-bioinfo.toulouse.inrae.fr). The DiCoExpress tool was used to analyze the RNAseq data (Lambert *et al*., 2020*a*). The differential expression analysis used generalized linear models. For the first experiment, we expressed the log of the average gene expression as a function of dpi. For the second experiment, we expressed the log of the average gene expression as an additive function of the effects of N treatment, dpi, and interaction between N treatment and dpi. For exp3, we performed two analyses separately at 2dpi and 7 dpi. In this case, we expressed the log of the average gene expression as an additive function of the effects of the N treatment, the genotype and the interaction between N treatment and the genotype. Likelihood ratio tests allowed to evaluate the expression changes associated to “dpi” in the first experiment, “N treatment” in exp2, to “N treatment” and interaction between “N treatment” and “genotype” in exp3. This test allowed to identified transcripts differentially regulated by N treatment in the sunn mutant as compared to the wild type in exp3. The Benjamini–Hochberg procedure allowed to adjust the probabilities of significance to control the false discovery rate (FDR). We used a thresholds of 0.05 to select differentially accumulated transcripts (DATs). We performed the co-expression analysis by using the ‘Coseq’ (v.1.4.0) algorithm (Rau and Maugis-Rabusseau, 2018) implemented and optimized in the DiCoexpress script (Lambert *et al*., 2020*a*). We complemented the MtV4 annotation with the Plant metabolic network (PMN)-Medicyc annotation of biochemical pathways (Urbanczyk-Wochniak and Sumner, 2007) to perform enrichment analysis using hypergeometric tests with the reference set defined as the whole genome. A functional enrichment was declared when the p-value of an annotation term was lower than 0.05. We performed targeted RT-qPCR on specific plant transcripts in a LightCycler (LightCycler 480; Roche Diagnostics) as previously described (Girin *et al*., 2007) using primers described in Supplemental table S13. The ACTIN11 and GAPDH A transcripts were used to normalize the data (Supplemental table S12).

## Acknowledgments

This work was supported by the ANR grants Psyché (ANR-16-CE20-0009) and LabEx Saclay Plant Sciences-SPS (ANR-10-LABX-0040-SPS). We thank Etienne-Pascal Journet for kindly providing seeds of A7/NL415 and sunn-2/NL415, Renaud Brouquisse and Pierre Frendo for critical reading of the manuscript.

## List of Author Contributions

M.L. designed the experiments; M.P. A.K. M.N. and M.L. performed the split-root experiments and the root developmental analysis; M.P., I.L., S.C., M.T., F.J., M-L.M-M. and M.L. performed the RNAseq analysis; M.P., A.K. M.N. performed RT-Q-PCR analysis; M.L. wrote the manuscript, with contributions of S.C and M.P. and revisions from all authors.

## References

Azarakhsh M, Lebedeva MA, Lutova LA. 2018. Identification and Expression Analysis of Medicago truncatula Isopentenyl Transferase Genes (IPTs) Involved in Local and Systemic Control of Nodulation. Frontiers in Plant Science 9, 304.

Bacanamwo M, Harper JE. 1997. The feedback mechanism of nitrate inhibition of nitrogenase activity in soybean may involve asparagine and/or products of its metabolism. Physiologia Plantarum 100, 371–377.

Bellegarde F, Gojon A, Martin A. 2017. Signals and players in the transcriptional regulation of root responses by local and systemic N signaling in Arabidopsis thaliana. Journal of Experimental Botany 68, 2553–2565.

van Brussel AAN, Tak T, Boot KJM, Kijne JW. 2002. Autoregulation of root nodule formation: signals of both symbiotic partners studied in a split-root system of Vicia sativa subsp. nigra. Molecular plant-microbe interactions: MPMI 15, 341–349.

Buhian WP, Bensmihen S. 2018. Mini-Review: Nod Factor Regulation of Phytohormone Signaling and Homeostasis During Rhizobia-Legume Symbiosis. Frontiers in Plant Science 9, 1247.

Cabeza R, Koester B, Liese R, Lingner A, Baumgarten V, Dirks J, Salinas-Riester G, Pommerenke C, Dittert K, Schulze J. 2014. An RNA sequencing transcriptome analysis reveals novel insights into molecular aspects of the nitrate impact on the nodule activity of Medicago truncatula. Plant Physiology 164, 400–411.

Chaput V, Martin A, Lejay L. 2020. Redox metabolism: the hidden player in carbon and nitrogen signaling? Journal of Experimental Botany 71, 3816–3826.

Charron D, Pingret J-L, Chabaud M, Journet E-P, Barker DG. 2004. Pharmacological Evidence That Multiple Phospholipid Signaling Pathways Link Rhizobium Nodulation Factor Perception in Medicago truncatula Root Hairs to Intracellular Responses, Including Ca2+ Spiking and Specific ENOD Gene Expression. Plant Physiology 136, 3582–3593.

Cho MJ, Harper JE. 1991. Effect of localized nitrate application on isoflavonoid concentration and nodulation in split-root systems of wild-type and nodulation-mutant soybean plants. Plant Physiology 95, 1106–1112.

Combier J-P, Vernié T, de Billy F, El Yahyaoui F, Mathis R, Gamas P. 2007. The MtMMPL1 early nodulin is a novel member of the matrix metalloendoproteinase family with a role in Medicago truncatula infection by Sinorhizobium meliloti. Plant Physiology 144, 703–716.

Drew MC. 1975. Comparison of the Effects of a Localised Supply of Phosphate, Nitrate, Ammonium and Potassium on the Growth of the Seminal Root System, and the Shoot, in Barley. New Phytologist 75, 479–490.

Ferguson BJ, Mathesius U. 2014. Phytohormone regulation of legume-rhizobia interactions. Journal of Chemical Ecology 40, 770–790.

Forde B, Lorenzo H. 2001. The nutritional control of root development. Plant and Soil 232, 51–68.

Gamas P, Brault M, Jardinaud M-F, Frugier F. 2017. Cytokinins in Symbiotic Nodulation: When, Where, What For? Trends in Plant Science 22, 792–802.

Gansel X, Muños S, Tillard P, Gojon A. 2001. Differential regulation of the NO3– and NH4+ transporter genes AtNrt2.1 and AtAmt1.1 in Arabidopsis: relation with long-distance and local controls by N status of the plant. The Plant Journal 26, 143–155.

Gautrat P, Laffont C, Frugier F. 2020. Compact Root Architecture 2 Promotes Root Competence for Nodulation through the miR2111 Systemic Effector. Current biology: CB 30, 1339-1345.e3.

Gent L, Forde BG. 2017. How do plants sense their nitrogen status? Journal of Experimental Botany 68, 2531–2539.

Girin T, Lejay L, Wirth J, Widiez T, Palenchar PM, Nazoa P, Touraine B, Gojon A, Lepetit M. 2007. Identification of a 150 bp cis-acting element of the AtNRT2.1 promoter involved in the regulation of gene expression by the N and C status of the plant. Plant, Cell & Environment 30, 1366–1380.

Goh C-H, Nicotra AB, Mathesius U. 2019. Genes controlling legume nodule numbers affect phenotypic plasticity responses to nitrogen in the presence and absence of rhizobia. Plant, Cell & Environment 42, 1747–1757.

Gonzalez-Rizzo S, Crespi M, Frugier F. 2006. The Medicago truncatula CRE1 Cytokinin Receptor Regulates Lateral Root Development and Early Symbiotic Interaction with Sinorhizobium meliloti. The Plant Cell 18, 2680–2693.

Guinel FC. 2015. Ethylene, a Hormone at the Center-Stage of Nodulation. Frontiers in Plant Science 6, 1121.

Hinson K. 1975. Nodulation Responses from Nitrogen Applied to Soybean Half-Root Systems1. Agronomy Journal 67, 799–804.

Huault E, Laffont C, Wen J, Mysore KS, Ratet P, Duc G, Frugier F. 2014. Local and systemic regulation of plant root system architecture and symbiotic nodulation by a receptor-like kinase. PLoS genetics 10, e1004891.

Jeudy C, Ruffel S, Freixes S, et al. 2010. Adaptation of Medicago truncatula to nitrogen limitation is modulated via local and systemic nodule developmental responses. The New Phytologist 185, 817–828.

Jia Z, von Wirén N. 2020. Signaling pathways underlying nitrogen-dependent changes in root system architecture: from model to crop species. Journal of Experimental Botany 71, 4393–4404.

Journet EP, El-Gachtouli N, Vernoud V, de Billy F, Pichon M, Dedieu A, Arnould C, Morandi D, Barker DG, Gianinazzi-Pearson V. 2001. Medicago truncatula ENOD11: a novel RPRP-encoding early nodulin gene expressed during mycorrhization in arbuscule-containing cells. Molecular plant-microbe interactions: MPMI 14, 737–748.

Kaló P, Gleason C, Edwards A, et al. 2005. Nodulation Signaling in Legumes Requires NSP2, a Member of the GRAS Family of Transcriptional Regulators. Science 308, 1786–1789.

Kassaw T, Bridges W, Frugoli J. 2015. Multiple Autoregulation of Nodulation (AON) Signals Identified through Split Root Analysis of Medicago truncatula sunn and rdn1 Mutants. Plants (Basel, Switzerland) 4, 209–224.

Kiers ET, Rousseau RA, West SA, Denison RF. 2003. Host sanctions and the legume-rhizobium mutualism. Nature 425, 78–81.

Kinkema M, Scott PT, Gresshoff PM. 2006. Legume nodulation: successful symbiosis through short- and long-distance signalling. Functional Plant Biology 33, 707–721.

Kinkema M, Gresshoff PM. 2008. Investigation of downstream signals of the soybean autoregulation of nodulation receptor kinase GmNARK. Molecular plant-microbe interactions: MPMI 21, 1337–1348.

Kosslak RM, Bohlool BB. 1984. Suppression of nodule development of one side of a split-root system of soybeans caused by prior inoculation of the other side. Plant Physiology 75, 125–130.

Krouk G, Lacombe B, Bielach A, et al. 2010. Nitrate-regulated auxin transport by NRT1.1 defines a mechanism for nutrient sensing in plants. Developmental Cell 18, 927–937.

Laffont C, Huault E, Gautrat P, Endre G, Kalo P, Bourion V, Duc G, Frugier F. 2019. Independent Regulation of Symbiotic Nodulation by the SUNN Negative and CRA2 Positive Systemic Pathways. Plant Physiology 180, 559–570.

Laffont C, Ivanovici A, Gautrat P, Brault M, Djordjevic MA, Frugier F. 2020. The NIN transcription factor coordinates CEP and CLE signaling peptides that regulate nodulation antagonistically. Nature Communications 11, 3167.

Lagarde D, Basset M, Lepetit M, Conejero G, Gaymard F, Astruc S, Grignon C. 1996. Tissue-specific expression of Arabidopsis AKT1 gene is consistent with a role in K+ nutrition. The Plant Journal: For Cell and Molecular Biology 9, 195–203.

Laguerre G, Heulin-Gotty K, Brunel B, Klonowska A, Le Quéré A, Tillard P, Prin Y, Cleyet-Marel J-C, Lepetit M. 2012. Local and systemic N signaling are involved in Medicago truncatula preference for the most efficient Sinorhizobium symbiotic partners. The New Phytologist 195, 437–449.

Lagunas B, Achom M, Bonyadi-Pour R, et al. 2019. Regulation of Resource Partitioning Coordinates Nitrogen and Rhizobia Responses and Autoregulation of Nodulation in Medicago truncatula. Molecular Plant 12, 833–846.

Lambert I, Paysant-Le Roux C, Colella S, Martin-Magniette M-L. 2020a. DiCoExpress: a tool to process multifactorial RNAseq experiments from quality controls to co-expression analysis through differential analysis based on contrasts inside GLM models. Plant Methods 16, 68.

Lambert I, Pervent M, Le Queré A, et al. 2020b. Responses of mature symbiotic nodules to the whole-plant systemic nitrogen signaling. Journal of Experimental Botany 71, 5039–5052.

Larrainzar E, Villar I, Rubio MC, Pérez-Rontomé C, Huertas R, Sato S, Mun J-H, Becana M. 2020. Hemoglobins in the legume–Rhizobium symbiosis. New Phytologist 228, 472–484.

Lejay L, Gansel X, Cerezo M, Tillard P, Müller C, Krapp A, von Wirén N, Daniel-Vedele F, Gojon A. 2003. Regulation of root ion transporters by photosynthesis: functional importance and relation with hexokinase. The Plant Cell 15, 2218–2232.

Lejay L, Wirth J, Pervent M, Cross JM-F, Tillard P, Gojon A. 2008. Oxidative pentose phosphate pathway-dependent sugar sensing as a mechanism for regulation of root ion transporters by photosynthesis. Plant Physiology 146, 2036–2053.

Li D, Kinkema M, Gresshoff PM. 2009. Autoregulation of nodulation (AON) in Pisum sativum (pea) involves signalling events associated with both nodule primordia development and nitrogen fixation. Journal of Plant Physiology 166, 955–967.

Li Y, Krouk G, Coruzzi GM, Ruffel S. 2014. Finding a nitrogen niche: a systems integration of local and systemic nitrogen signalling in plants. Journal of Experimental Botany 65, 5601–5610.

Marsh JF, Rakocevic A, Mitra RM, Brocard L, Sun J, Eschstruth A, Long SR, Schultze M, Ratet P, Oldroyd GED. 2007. Medicago truncatula NIN Is Essential for Rhizobial-Independent Nodule Organogenesis Induced by Autoactive Calcium/Calmodulin-Dependent Protein Kinase. Plant Physiology 144, 324–335.

Mens C, Hastwell AH, Su H, Gresshoff PM, Mathesius U, Ferguson BJ. 2020. Characterisation of Medicago truncatula CLE34 and CLE35 in nitrate and rhizobia regulation of nodulation. The New Phytologist.

Mergaert P, Kereszt A, Kondorosi E. 2020. Gene Expression in Nitrogen-Fixing Symbiotic Nodule Cells in Medicago truncatula and Other Nodulating Plants. The Plant Cell 32, 42–68.

Middleton PH, Jakab J, Penmetsa RV, et al. 2007. An ERF Transcription Factor in Medicago truncatula That Is Essential for Nod Factor Signal Transduction. The Plant Cell 19, 1221–1234.

Mohd-Radzman NA, Laffont C, Ivanovici A, Patel N, Reid D, Stougaard J, Frugier F, Imin N, Djordjevic MA. 2016. Different Pathways Act Downstream of the CEP Peptide Receptor CRA2 to Regulate Lateral Root and Nodule Development. Plant Physiology 171, 2536–2548.

Moreau S, Verdenaud M, Ott T, Letort S, de Billy F, Niebel A, Gouzy J, de Carvalho-Niebel F, Gamas P. 2011. Transcription reprogramming during root nodule development in Medicago truncatula. PloS One 6, e16463.

Mortier V, De Wever E, Vuylsteke M, Holsters M, Goormachtig S. 2012a. Nodule numbers are governed by interaction between CLE peptides and cytokinin signaling. The Plant Journal: For Cell and Molecular Biology 70, 367–376.

Mortier V, Holsters M, Goormachtig S. 2012b. Never too many? How legumes control nodule numbers. Plant, Cell & Environment 35, 245–258.

Nakagawa T, Kawaguchi M. 2006. Shoot-applied MeJA suppresses root nodulation in Lotus japonicus. Plant & Cell Physiology 47, 176–180.

Nishida H, Suzaki T. 2018. Two Negative Regulatory Systems of Root Nodule Symbiosis: How Are Symbiotic Benefits and Costs Balanced? Plant & Cell Physiology 59, 1733–1738.

Nishida H, Tanaka S, Handa Y, et al. 2018. A NIN-LIKE PROTEIN mediates nitrate-induced control of root nodule symbiosis in Lotus japonicus. Nature Communications 9, 499.

Ohkubo Y, Tanaka M, Tabata R, Ogawa-Ohnishi M, Matsubayashi Y. 2017. Shoot-to-root mobile polypeptides involved in systemic regulation of nitrogen acquisition. Nature Plants 3, 17029.

Okamoto S, Shinohara H, Mori T, Matsubayashi Y, Kawaguchi M. 2013. Root-derived CLE glycopeptides control nodulation by direct binding to HAR1 receptor kinase. Nature Communications 4, 2191.

Okamoto S, Tabata R, Matsubayashi Y. 2016. Long-distance peptide signaling essential for nutrient homeostasis in plants. Current Opinion in Plant Biology 34, 35–40.

Oldroyd GED, Murray JD, Poole PS, Downie JA. 2011. The rules of engagement in the legume-rhizobial symbiosis. Annual Review of Genetics 45, 119–144.

Omrane S, Ferrarini A, D’Apuzzo E, Rogato A, Delledonne M, Chiurazzi M. 2009. Symbiotic competence in Lotus japonicus is affected by plant nitrogen status: transcriptomic identification of genes affected by a new signalling pathway. The New Phytologist 183, 380–394.

Ota R, Ohkubo Y, Yamashita Y, Ogawa-Ohnishi M, Matsubayashi Y. 2020. Shoot-to-root mobile CEPD-like 2 integrates shoot nitrogen status to systemically regulate nitrate uptake in Arabidopsis. Nature Communications 11.

Ott T, van Dongen JT, Günther C, Krusell L, Desbrosses G, Vigeolas H, Bock V, Czechowski T, Geigenberger P, Udvardi MK. 2005. Symbiotic leghemoglobins are crucial for nitrogen fixation in legume root nodules but not for general plant growth and development. Current biology: CB 15, 531–535.

Pecrix Y, Staton SE, Sallet E, et al. 2018. Whole-genome landscape of Medicago truncatula symbiotic genes. Nature Plants 4, 1017–1025.

Pérez Guerra JC, Coussens G, De Keyser A, De Rycke R, De Bodt S, Van De Velde W, Goormachtig S, Holsters M. 2010. Comparison of developmental and stress-induced nodule senescence in Medicago truncatula. Plant Physiology 152, 1574–1584.

Prell J, White JP, Bourdes A, Bunnewell S, Bongaerts RJ, Poole PS. 2009. Legumes regulate Rhizobium bacteroid development and persistence by the supply of branched-chain amino acids. Proceedings of the National Academy of Sciences of the United States of America 106, 12477–12482.

Rau A, Maugis-Rabusseau C. 2018. Transformation and model choice for RNA-seq co-expression analysis. Briefings in Bioinformatics 19, 425–436.

Reid DE, Ferguson BJ, Gresshoff PM. 2011a. Inoculation- and nitrate-induced CLE peptides of soybean control NARK-dependent nodule formation. Molecular plant-microbe interactions: MPMI 24, 606–618.

Reid DE, Ferguson BJ, Hayashi S, Lin Y-H, Gresshoff PM. 2011b. Molecular mechanisms controlling legume autoregulation of nodulation. Annals of Botany 108, 789–795.

Roux B, Rodde N, Jardinaud M-F, et al. 2014. An integrated analysis of plant and bacterial gene expression in symbiotic root nodules using laser-capture microdissection coupled to RNA sequencing. The Plant Journal: For Cell and Molecular Biology 77, 817–837.

Ruffel S, Freixes S, Balzergue S, et al. 2008. Systemic signaling of the plant nitrogen status triggers specific transcriptome responses depending on the nitrogen source in Medicago truncatula. Plant Physiology 146, 2020–2035.

Ruffel S, Gojon A, Lejay L. 2014. Signal interactions in the regulation of root nitrate uptake. Journal of Experimental Botany 65, 5509–5517.

Sagan M, Morandi D, Tarenghi E, Duc G. 1995. Selection of nodulation and mycorrhizal mutants in the model plant Medicago truncatula (Gaertn.) after γ-ray mutagenesis. Plant Science 111, 63–71.

Sasaki T, Suzaki T, Soyano T, Kojima M, Sakakibara H, Kawaguchi M. 2014. Shoot-derived cytokinins systemically regulate root nodulation. Nature Communications 5, 4983.

Schnabel E, Journet E-P, de Carvalho-Niebel F, Duc G, Frugoli J. 2005. The Medicago truncatula SUNN Gene Encodes a CLV1-likeLeucine-rich Repeat Receptor Kinase that Regulates Nodule Number and Root Length. Plant Molecular Biology 58, 809–822.

Seabra AR, Pereira PA, Becker JD, Carvalho HG. 2012. Inhibition of glutamine synthetase by phosphinothricin leads to transcriptome reprograming in root nodules of Medicago truncatula. Molecular plant-microbe interactions: MPMI 25, 976–992.

Silsbury JH, Catchpoole DW, Wallace W. 1986. Effects of Nitrate and Ammonium on Nitrogenase (C2H2 Reduction) Activity of Swards of Subterranean Clover, Trifolium subterraneum L. Functional Plant Biology 13, 257–273.

Stitt M, Müller C, Matt P, Gibon Y, Carillo P, Morcuende R, Scheible W-R, Krapp A. 2002. Steps towards an integrated view of nitrogen metabolism. Journal of Experimental Botany 53, 959–970.

Tabata R, Sumida K, Yoshii T, Ohyama K, Shinohara H, Matsubayashi Y. 2014. Perception of root-derived peptides by shoot LRR-RKs mediates systemic N-demand signaling. Science (New York, N.Y.) 346, 343–346.

Tanaka A, Fujlta K, Terasawa H. 1985. Growth and Dinitrogen Fixation, of Soybean Root System Affected by Partial Exposure to Nitrate. Soil Science and Plant Nutrition 31, 637–645.

Tsikou D, Yan Z, Holt DB, Abel NB, Reid DE, Madsen LH, Bhasin H, Sexauer M, Stougaard J, Markmann K. 2018. Systemic control of legume susceptibility to rhizobial infection by a mobile microRNA. Science (New York, N.Y.) 362, 233–236.

Urbanczyk-Wochniak E, Sumner LW. 2007. MedicCyc: a biochemical pathway database for Medicago truncatula. Bioinformatics 23, 1418–1423.

Valkov VT, Rogato A, Alves LM, Sol S, Noguero M, Léran S, Lacombe B, Chiurazzi M. 2017. The Nitrate Transporter Family Protein LjNPF8.6 Controls the N-Fixing Nodule Activity. Plant Physiology 175, 1269–1282.

Vidal EA, Alvarez JM, Araus V, et al. 2020. Nitrate in 2020: Thirty Years from Transport to Signaling Networks. The Plant Cell 32, 2094–2119.

Xia X, Ma C, Dong S, Xu Y, Gong Z. 2017. Effects of nitrogen concentrations on nodulation and nitrogenase activity in dual root systems of soybean plants. Soil Science and Plant Nutrition 63, 470–482.

